# Targeting PIEZO1-TMEM16F Coupling to Mitigate Sickle Cell Disease Complications

**DOI:** 10.1101/2025.05.27.656389

**Authors:** Pengfei Liang, Yui-Chun Serena Wan, Ke Zoe Shan, Ryan Chou, Yang Zhang, Martha Delahunty, Sanjay Khandelwal, Samuel J. Francis, Gowthami M. Arepally, Marilyn J. Telen, Huanghe Yang

## Abstract

A deeper understanding of sickle cell disease (SCD) pathophysiology is critical for identifying novel therapeutic targets. A hallmark of SCD is abnormal phosphatidylserine (PS) exposure on sickle red blood cells (RBCs), which contributes to anemia, thrombosis, and vaso-occlusive crises (VOC). However, the mechanisms underlying this excessive PS exposure remain unclear. Here, we identify TMEM16F, a Ca^2+^-activated lipid scramblase, as a key mediator of PS exposure downstream of Ca^2+^ influx through the mechanosensitive channel PIEZO1 in sickle RBCs. Electrophysiology, imaging and flow cytometry reveal that deoxygenation-induced sickling promotes PIEZO1 activation, triggering Ca^2+^ entry, TMEM16F activation, and PS exposure. This cascade enhances PS^+^ microparticle release, thrombin generation, and RBC adhesion to endothelial cells. Notably, partial PIEZO1 inhibition with benzbromarone, an anti-gout drug, suppresses these changes. Our findings thus define a previously unrecognized mechanotransduction pathway in sickle RBCs and propose a unique therapeutic strategy to mitigate hypercoagulability and vaso-occlusion associated with SCD.

**Brief Summary:** Enhanced PIEZO1 activation in sickle red blood cells promotes TMEM16F scramblase-mediated phosphatidylserine exposure and subsequent sickle cell disease complications. Disrupting this coupling presents a potential therapeutic strategy.

## Introduction

Sickle cell disease (SCD) is the most common inherited blood disorder that affects millions of people worldwide, including ∼100,000 individuals in the United States and 1 in 500 African Americans (Hassell, 2010). Despite advances in chemical, cellular, and gene-based therapies, SCD remains a significant global public health challenge (Migotsky et al., 2022; Piel et al., 2023; Telen, 2020). In order to develop next generation therapeutics to treat SCD, identification of new therapeutic targets is urgently needed.

It is well-documented that sickle RBCs (SS RBCs) exhibit elevated exposure of PS (Wood et al., 1996; Yasin et al., 2003), a phospholipid normally confined to the inner leaflet of the plasma membrane. This aberrant PS exposure contributes to several major complications of SCD, including anemia, hypercoagulability, and VOC. On one hand, PS serves as an “eat-me” signal to mediate erythrophagocytosis of SS RBCs, thereby exacerbating anemia (de Back et al., 2014). On the other hand, both PS-exposed (PS^+^) SS RBCs and PS^+^ RBC-derived microparticles (RMPs) collectively expand the procoagulant surface area, driving a hypercoagulable state (Ataga and Key, 2007; Ataga and Orringer, 2003; Garnier et al., 2020). In addition, PS exposure enhances the abnormal adhesion of SS RBCs to endothelial cells, a key event in VOC that triggers pain crises and progressive organ damage (Kaul et al., 2009b). Therefore, PS exposure is a critical contributor to SCD pathophysiology and a compelling therapeutic target. Despite its clinical importance, the molecular mechanisms responsible for enhanced PS exposure in SS RBCs remain poorly understood, posing a barrier to the development of targeted interventions.

We demonstrated that TMEM16F is the sole Ca^2+^-activated phospholipid scramblase (CaPLSase) in RBCs (Liang et al., 2024; Yang et al., 2012). TMEM16F senses Ca^2+^ influx through PIEZO1, a mechanosensitive cation channel in RBCs (Cahalan et al., 2015; Vaisey et al., 2022; Zarychanski et al., 2012), and drives rapid PS exposure (Liang et al., 2024). Notably, the functional coupling between PIEZO1 and TMEM16F is markedly enhanced in RBCs from patients with hereditary xerocytosis (HX) patients carrying PIEZO1 gain-of-function (GOF) mutations, highlighting the importance of this signaling axis in red cell pathophysiology (Liang et al., 2024). Elevated intracellular Ca²⁺ is a well-recognized trigger of PS exposure in SS RBCs (Weiss et al., 2012; Weiss et al., 2011), and recent evidence implicates PIEZO1 as a major source of increased Ca²⁺ permeability in these cells (Nader et al., 2023; Wadud et al., 2020). Stimulation of PIEZO1 by its agonist Yoda1 increases Ca²⁺ influx, which promotes RBC dehydration, exacerbates sickling, augments PS exposure, and enhances SS RBC adhesion to endothelial cells (Nader et al., 2023). Taken together, these findings support a model in which hyperactivation of the PIEZO1-TMEM16F axis drives excessive PS exposure in SS RBCs and contributes to the hypercoagulability and increased vascular adhesion observed in sickle cell disease (SCD).

To test this hypothesis, we systematically investigated intracellular Ca²⁺ dynamics, PS exposure, RMP release, prothrombinase activity, and RBC-endothelium adhesion in SS RBCs. Our findings demonstrate that deoxygenation-induced sickling triggers Ca²⁺ influx through PIEZO1, which in turn activates TMEM16F, leading to increased PS exposure, RMP release, and thrombin generation. Moreover, enhanced coupling between PIEZO1 and TMEM16F promotes the adhesion of SS RBCs to endothelial cells. Importantly, pharmacological inhibition of PIEZO1 effectively disrupts this pathogenic cascade, preventing sickling-induced PS externalization, microparticle release, thrombin generation, and endothelial adhesion. These results uncover a critical role for the PIEZO1-TMEM16F axis in SCD-associated hypercoagulability and VOC, and suggest a novel therapeutic strategy to mitigate these complications.

## Results

### Enhanced PIEZO1 activity contributes to Ca^2+^ elevation in sickled RBCs

Recent studies reported that PIEZO1 contributes to intracellular Ca^2+^ ([Ca^2+^]i) increase in SS RBCs (Nader et al., 2023; Wadud et al., 2020), a well-documented phenomenon since it was first reported in 1973 (Eaton et al., 1973). Our Fura-2 measurement confirmed this observation (Figure 1A). SS RBCs have significantly higher basal [Ca^2+^]i than healthy donor RBCs (AA

**Figure 1.**
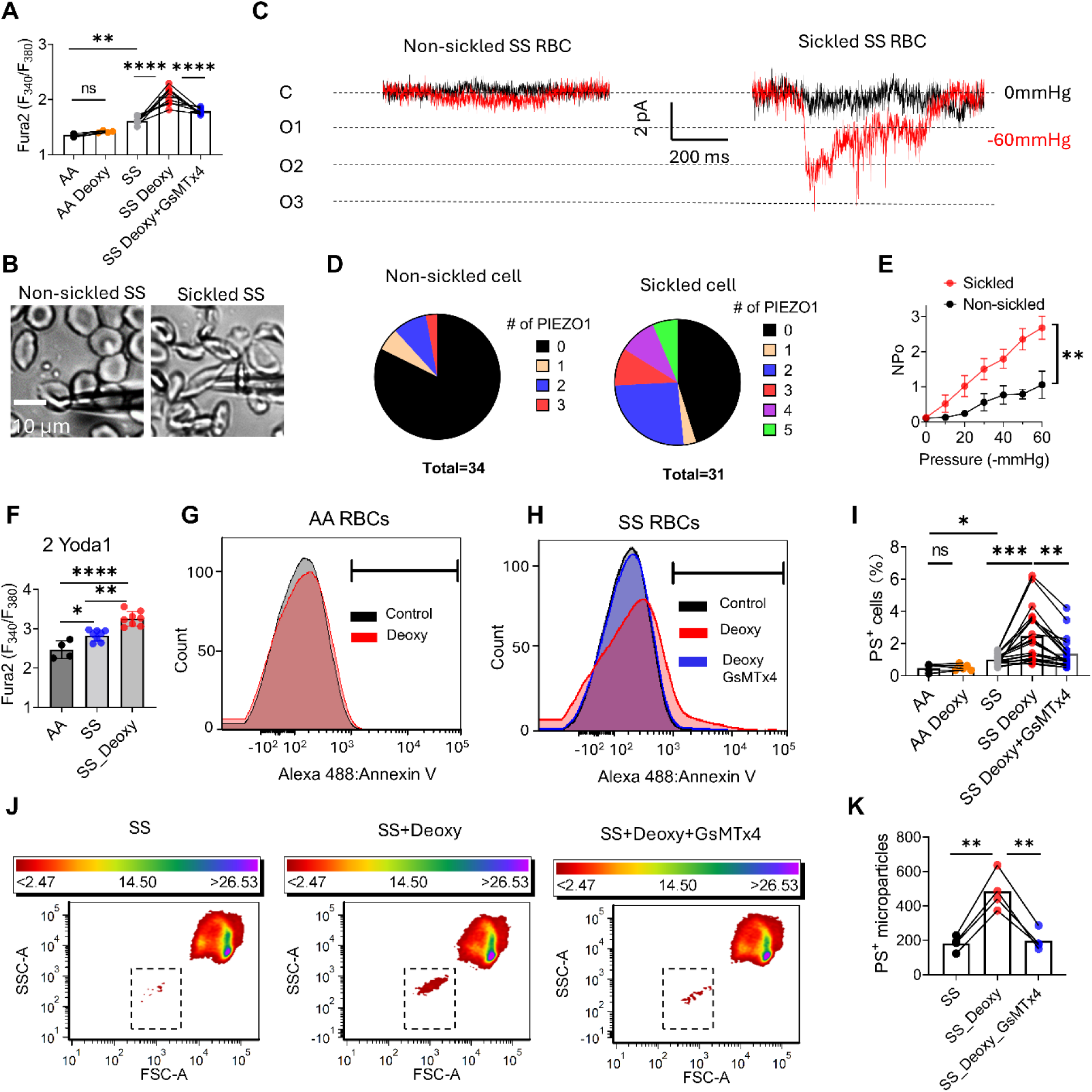
Enhanced PIEZO1 activity contributes to Ca^2+^ increase, phosphatidylserine (PS) exposure and microparticle release in deoxygenated SS RBCs. (A) Basal intracellular Ca²⁺ levels in red blood cells (RBCs) from healthy donors (AA) and sickle cell disease patients (SS) under normoxic condition and deoxygenated (Deoxy) conditions with and without 5 µM GsMTx4, a PIEZO1 inhibitor. One-way ANOVA followed by Tukey’s test. **P < 0.01, ****P < 0.0001, ns: no significance. Sample sizes: n = 4 for healthy donors and n = 8 for sickle cell patients. Data are presented as the means of triplicates for each sample. (B) Representative images of press-clamp measurement on non-sickled and sickled SS RBCs. (C) Representative mechanosensitive currents recorded from non-sickled and sickled SS RBCs. Black traces: no current was elicited at 0 mmHg (C: closed state); Red traces: currents were recorded at -60 mmHg pressure. Multiple open states (O1, O2, O3) were frequently observed in sickled SS RBCs. (D) Quantification of the number of the mechanosensitive channels recorded in single patches from non-sickled (n=34) and sickled (n=31) SS RBCs. (E) The NPo (number of channels multiplied by the open probability) of the mechanosensitive channel in non-sickled and sickled SS RBCs elicited by various negative pressures. Unpaired two-sided Student’s t-test (**P < .01). Sample sizes: n = 34 for non-sickled and n = 31 for sickled SS RBCs. (F) 2 µM Yoda1-induced Ca²⁺ increase in RBCs from healthy donors (AA) and SCD patients (SS) under normoxic and deoxygenated (Deoxy) conditions. The data represents the peak of the Ca²⁺ levels. One-way ANOVA followed by Tukey’s test. *P < 0.05, **P < 0.01, ****P < 0.0001. Sample sizes: n = 4 for AA and n = 6 for SS. Data are presented as the means of triplicates for each sample. (G) Flow cytometry analysis of PS exposure in RBCs from healthy donors (AA) under normoxic and deoxygenated conditions. (H) Flow cytometry analysis of PS exposure in SS RBCs under normoxic and deoxygenated conditions, with or without 5 µM GsMTx4. (I) Percentage of PS-positive cells under the conditions described in G and H. One-way ANOVA followed by Tukey’s test. *P < 0.05, **P < 0.01, ***P < .001, ns: no significance. Sample sizes: n = 5 for healthy donors and n = 20 for sickle cell patients. Data are presented as the means of triplicates for each sample. (J) Representative flow cytometry plots of microparticle release (dotted rectangles) in SS RBCs under three different conditions: control (left), deoxygenation (middle), and deoxygenation with 5 µM GsMTx4 (right). (K) Quantification of PS-positive microparticles under the conditions described in J. One-way ANOVA followed by Tukey’s test. ****P < 0.0001. Data were obtained from 4 SCD patients with triplicates for each sample. The lines in panels A, I and K connect data from the same individuals.

RBCs). Interestingly, deoxygenation induces a significant increase of basal [Ca^2+^]i, which can be prevented by GsMTx4, a PIEZO1 inhibitor (Bae et al., 2011), in SS RBCs (Figure 1A). Such deoxygenation-induced Ca^2+^ increase was not observed in AA RBCs, suggesting that deoxygenation-induced irreversible sickling, but not oxidative stress per se may promote PIEZO1 activation.

To further dissect, we employed the pressure-clamp technique to quantify PIEZO1 channel activity in SS RBCs by applying negative pressure (-60 mmHg) to the recorded membrane, thereby mechanically activating PIEZO1(Vaisey et al., 2022) (Figure 1B-C). We detected mechanosensitive single-channel activity with a conductance of 27.5 pS (Figure S1), consistent with the characteristic unitary conductance of the PIEZO1 channel (Coste et al., 2010). Our pressure clamp recording and single channel analysis further demonstrated a low probability (6 out of 34 patches) of detecting PIEZO1 current in non-sickled SS RBCs (Figure 1D), consistent with the low expression level of PIEZO1 (∼100 channels per RBC) (Vaisey et al., 2022). In the membrane patches with mechanosensitive current, no more than three PIEZO1 channels can be simultaneously recorded. In stark contrast, 14 out of 31 patches from sickled SS RBCs gave rise to 1-5 PIEZO1 channels, suggesting that PIEZO1 has a much higher surface density in sickled than non-sickled SS RBCs. Furthermore, in the membrane patches with detectable PIEZO1 activity, the open probability of PIEZO1 was significantly higher in sickled cells compared to non-sickled cells (Figure 1E), supporting that PIEZO1 channels become more active after sickling. To further validate, we examined the effect of Yoda1, a specific PIEZO1 agonist (Syeda et al., 2015), on [Ca^2+^]i. We found that Yoda1 induced a significantly higher [Ca^2+^]i in SS RBCs under normoxic condition, which was further exacerbated by deoxygenation (Figure 1F).

Taken together, our Ca^2+^ imaging and electrophysiology results demonstrate that PIEZO1 becomes more active after deoxygenation-induced irreversible sickling, contributing to elevated [Ca^2+^]i.

### PIEZO1 contributes to sickling-induced PS exposure and PS^+^ microparticle release

Next, we investigated if enhanced PIEZO1-mediated Ca^2+^ influx after sickling is correlated with elevated PS exposure and PS^+^ RBCs microparticle release (RMP), two common features of SS RBCs (Wood et al., 1996; Yasin et al., 2003). Our flow cytometry analysis using the PS probe AnV confirmed that SS RBCs have a significantly higher percentage of PS^+^ cells than AA RBCs (1.1% vs 0.4%, Figure 1G-I) under basal conditions. Deoxygenation markedly increased the PS^+^ population to 2.5% in SS samples (Figure 1H) without affecting PS exposure in AA RBCs (Figure 1G). GsMTx4 prevents this deoxygenation-induced PS exposure in SS RBCs, maintaining the PS^+^ population at 1.4% (Fig.1I). In addition, deoxygenation significantly enhanced PS^+^ RMP release from SS RBCs, which was also prevented by GsMTx4 (Figure 1J-K). Our flow cytometry experiments thus demonstrated that deoxygenation could increase basal PS exposure in SS RBCs and this increase is correlated with sickling-induced PIEZO1 activation (Figure 1A-E).

Then, we used 2 µM Yoda1 to activate PIEZO1 to mimic the mechanical stimulation of SS RBCs (Figure 2) (Nader et al., 2023). Our flow cytometry analysis showed that deoxygenation dramatically promotes Yoda1-induced PS exposure in SS RBCs as evidenced by a significant increase of PS^+^ population from 40% to 70% (15 SCD patients, Figure 2C-D). On the contrary, deoxygenation has no obvious effect on Yoda1-induced PS exposure in AA RBCs (Figure 2A-B). Confocal fluorescence microscopy at single-cell resolution further supports our observation using flow cytometry. Deoxygenated SS RBCs showed much faster, stronger, and more complete Yoda1-induced PS exposure than before deoxygenation (Figure 2E-G and Supplementary Videos 1-2). None of the cells underwent hemolysis under this condition, indicating that the observed PS exposure is not due to loss of membrane integrity. It is worth noting that the sickled SS RBCs showed more robust Yoda1-induced PS exposure than the non-sickled, biconcave-shaped SS RBCs (Figures. 2E-F and S2), further supporting that sickling promotes PIEZO1 activation and PS exposure. In addition, deoxygenation also significantly increased Yoda1-induced PS^+^ RMPs release from SS RBCs (Figure 2H-I). Our flow cytometry and imaging experiments thus demonstrate that deoxygenation-induced sickling greatly promotes PIEZO1-mediated PS exposure and PS^+^ RMP release.

**Figure 2.**
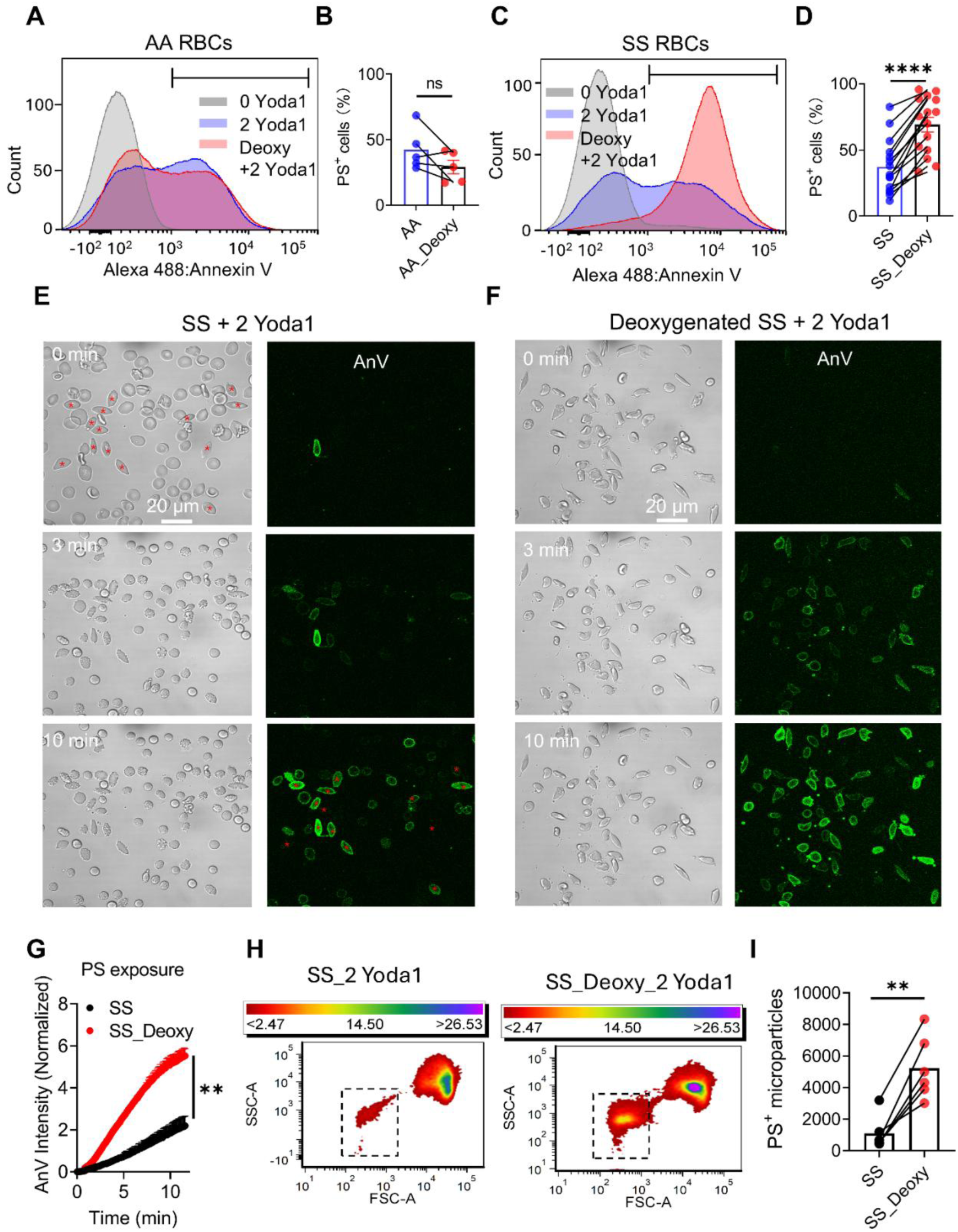
Sickling enhances Yoda1-induced PS exposure and microparticle release. (A) Representative flow cytometry analysis of PS exposure in RBCs from AA under normoxic (blue) and deoxygenated (red) conditions in the presence of 2 µM Yoda1. 0 Yoda1 under normoxic conditions (grey) was shown as a control. (B) Effect of deoxygenation on Yoda1-induced PS exposure in AA RBCs. Unpaired two-sided Student’s t-test. ns: no significance. n=5 healthy donors. Data are presented as the mean of triplicate for each sample. (C) Representative flow cytometry analysis of PS exposure in SS RBCs under normoxic (blue) and deoxygenated (red) conditions in the presence of 2 µM Yoda1. 0 Yoda1 under normoxic conditions (grey) was shown as a control. (D) Summary of 2 µM Yoda1-induced PS exposure in SS RBCs with and without deoxygenation. Unpaired two-sided Student’s t-test. ****P < 0.0001. n=15 SCD patients. Data are presented as the means of triplicate for one sample. (E-F) Representative confocal images of 2 µM Yoda1-induced PS exposure reported by Annexin V (AnV) in SS RBCs under normoxic (E) and deoxygenated (F) conditions, respectively. The sickled cells in E were labeled with red stars. (G) Time course of Yoda1-induced PS exposure in SS RBCs under normoxic and deoxygenated conditions. Unpaired two-sided Student’s t-test at the last time point. **P < 0.01. Data are presented as the means of triplicate for one sample. (H) Representative flow cytometry plots of Yoda1-induced microparticle release (dotted rectangles) from SS RBCs under normoxic (left) and deoxygenated (right) conditions. (I) Quantification of PS^+^ microparticles under the conditions described in (H). Unpaired two-sided Student’s t-test. ****P < 0.0001. Data were obtained from 6 SCD patients, with triplicate for each sample. The lines in panels B, D and I connect data from the same individuals.

### PIEZO1-TMEM16F coupling is augmented in sickled RBCs

The removal of extracellular Ca²⁺ abolished deoxygenation-induced PS exposure and the release of PS⁺ RMPs (Figure S3), confirming that these processes are Ca²⁺ dependent (Nader et al., 2023). We identified TMEM16F as the sole CaPLSase in RBCs, which spatially stays in the proximity of PIEZO1 and functionally senses PIEZO1-mediated Ca²⁺ influx (Liang et al., 2024; Yang et al., 2012). TMEM16F is also known to play an essential role in PS^+^ MP release from platelets (Fujii et al., 2015). Based on these findings, we hypothesize that sickling-induced PIEZO1 activation may enhance PIEZO1-TMEM16F coupling in SS RBCs and promote PS exposure and PS^+^ RMP release.

We first tested this hypothesis by utilizing TMEM16F’s moonlighting function as an ion channel (Liang and Yang, 2021; Yang et al., 2012; Zhang et al., 2022). Through cell-attached patch-clamp recordings, we quantify PIEZO1-TMEM16F coupling in response to Yoda1 stimulation (Figure 3). Under this condition, Yoda1-induced Ca^2+^ influx through PIEZO1 activates TMEM16F current (Figure 3A, left). Using Cs^+^ to inhibit Gardos Ca^2+^-activated K^+^ (KCa3.1) channels (see Methods) and performing current subtraction, we isolated the typical time- and voltage-dependent TMEM16F current (Liang and Yang, 2021; Liang et al., 2024; Yang et al., 2012; Zhang et al., 2022) (Figure 3A, right). The removal of extracellular Ca^2+^ completely abolished the Yoda1-induced current (Figure 3A, bottom), further supporting that the recorded current is Ca^2+^-dependent TMEM16F current. Our quantification showed that Yoda1-stimulated TMEM16F current in sickled SS RBCs (87.84 ± 7.58 pA) is significantly larger than the currents in AA RBCs (48.79 ± 5.01 pA) and non-sickled SS RBCs (48.98 ± 2.10 pA) at +160 mV (Figure 3B-C), further demonstrating that the functional coupling between PIEZO1 and TMEM16F is enhanced in sickled SS RBCs.

**Figure 3.**
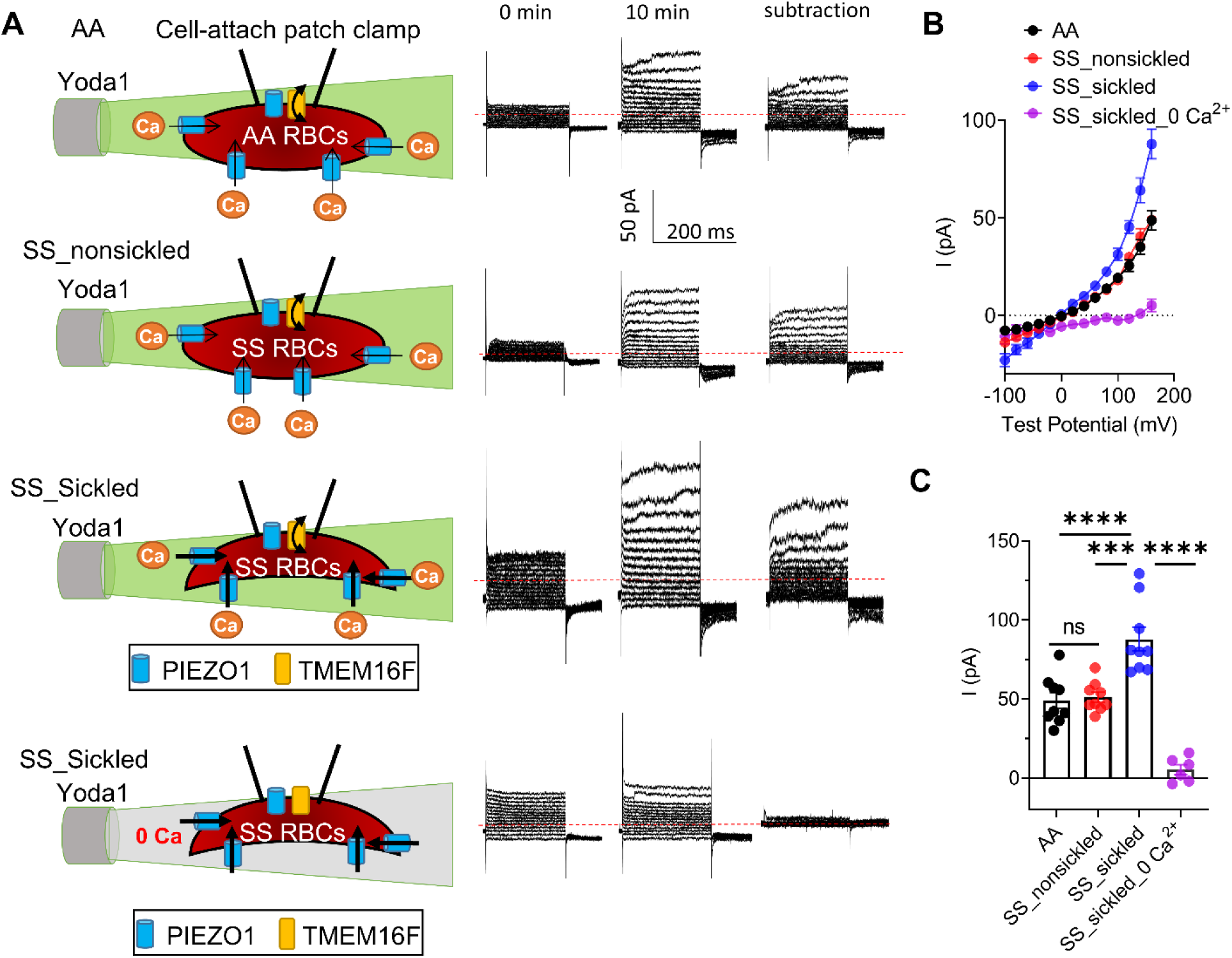
PIEZO1-TMEM16F coupling is augmented in sickled SS RBCs. (A) TMEM16F current elicited by 2 µM Yoda1 under four different conditions: healthy donor (AA) RBCs, non-sickled SS RBCs, sickled SS RBCs, and sickled SS RBCs with 0 extracellular Ca^2+^ (top to bottom). Voltage steps ranging from –100 mV to +160 mV were applied in 20 mV increments with a holding potential of –60 mV. Left: schematics of the cell-attached patch clamp conditions utilizing PIEZO1-mediated Ca^2+^ to activate TMEM16F within the same patch membrane. Right: representative currents before Yoda1 (left), 10 minutes after Yoda1 application (middle), and TMEM16F current derived by subtracting current before and after Yoda1 (right). (B) Current-voltage (I-V) relationships in A. Data are presented as mean ± SEM. n=6-9. (C) Yoda1-induced TMEM16F current amplitudes at +160 mV. Data are presented as mean ± SEM. n=6-9. One-way ANOVA followed by Tukey’s test. ***P < 0.001, ****P < 0.0001, ns: no significance.

### PIEZO1-TMEM16F coupling mediates RMP release

To establish the role of PIEZO1-TMEM16F coupling in RMP release, we stimulated the RBCs from the wild-type (WT) and TMEM16F knockout (KO) mice (Liang et al., 2024; Yang et al., 2012) with various concentrations of Yoda1 (Figure 4). Our flow cytometry analysis demonstrated that Yoda1 dose-dependently promotes PS^+^ RMP release and TMEM16F deficiency largely diminished Yoda1-induced PS^+^ RMP shedding. Our experiments thus showed that TMEM16F is required for PS^+^ RMPs shedding, and PIEZO1-TMEM16F coupling is sufficient to trigger PS^+^ RMPs release. Given the increased PIEZO1 activity and [Ca^2+^]i in sickled SS RBCs (Figure 1), the augmented PS^+^ RMPs release from these cells (Figures 1J-K and 2H-I) is likely due to enhanced PIEZO1-TMEM16F coupling.

**Figure 4.**
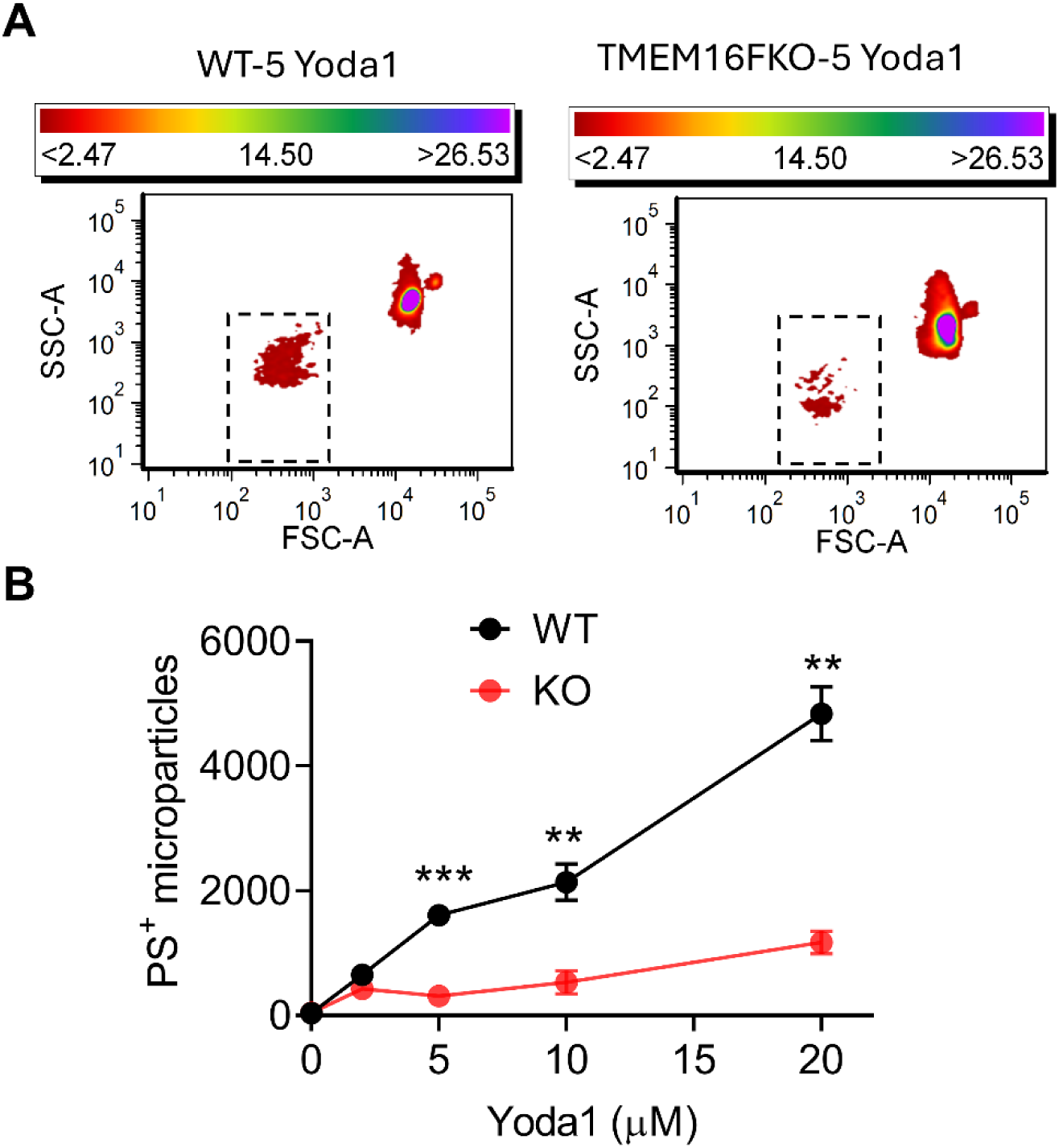
TMEM16F controls Yoda1-induced PS^+^ microparticle release from RBCs. (A) Representative flow cytometry plots of 5 µM Yoda1-induced microparticle release (dotted rectangles) from WT and TMEM16FKO mice RBCs. (B) Quantification of Yoda1-induced PS^+^ microparticle release from the RBCs from wild-type (WT) and TMEM16F-knockout (KO) mice. Two-way ANOVA followed by Tukey’s test. **P<0.01, ***P < 0.001. Data were obtained from 2 WT and 2 KO mice, with triplicates for each sample.

### GsMTx-4 suppresses deoxygenation-induced PS exposure and prothrombinase activity in SS RBCs

Having identified the molecular mechanism for enhanced PS exposure and PS^+^ RMPs release in SS RBCs, we next investigated if disrupting PIEZO1-TMEM16F coupling could be a feasible strategy to prevent lipid scrambling-mediated SCD complications, especially after sickling. As we have shown that GsMTx-4 suppresses basal [Ca^2+^]i elevation PS exposure and PS^+^ RMPs release in SS RBCs (Figure 1A, 1I and 1K), we conducted proof-of-principle experiments to test whether decoupling PIEZO1-TMEM16F with the PIEZO1 inhibitor could mitigate deoxygenation-induced changes.

GsMTx-4 dose-dependently suppressed Yoda1-induced Ca²⁺ influx (IC₅₀ = 3.66 ± 0.72 μM, Figure 5A, C-D) and PS exposure (IC₅₀ = 1.27 ± 0.12 μM, Figure 5B, C-D) in deoxygenated SS RBCs. Notably, the IC₅₀ for inhibiting PS exposure was significantly lower than that for suppressing Ca²⁺ influx (Figure 5C-D), suggesting there is a therapeutic window to completely prevent PS exposure in SS RBCs without completely inhibiting PIEZO1. We previously also observed a similar therapeutic window when targeting PIEZO-TMEM16F coupling in HX RBCs (Liang et al., 2024). Therefore, partial inhibition of PIEZO1 can effectively decouple PIEZO-TMEM16F functional interaction in sickled RBCs while maintaining PIEZO1’s essential functions in RBCs and other cell types.

**Figure 5.**
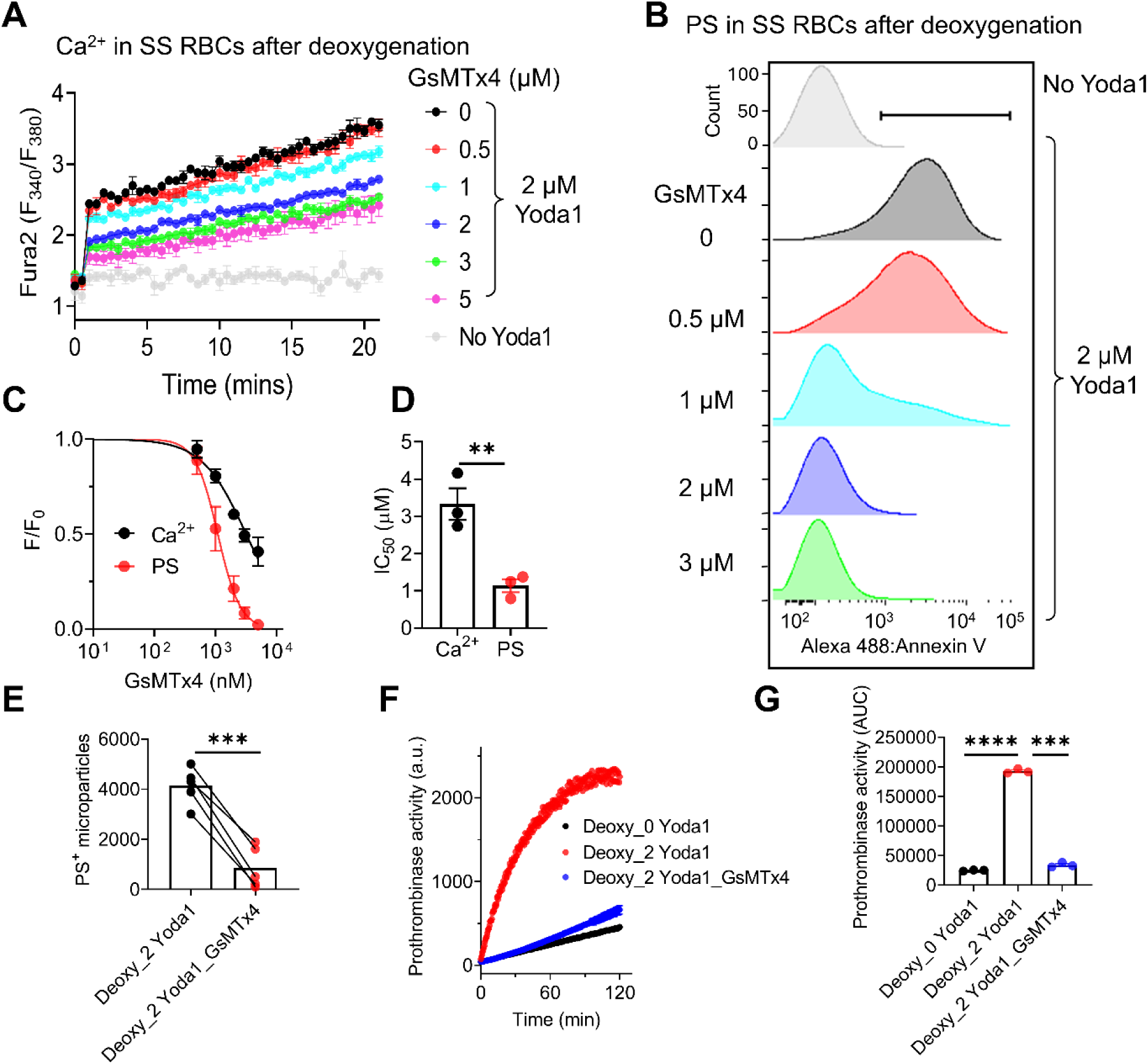
GsMTx4 suppresses Yoda1-induced Ca^2+^ increase, PS exposure and thrombin generation in deoxygenated SS RBCs. (A) Fura2 measurements of Yoda1-induced Ca²⁺ influx in deoxygenated SS RBCs in response to different concentrations of GsMTx-4. Data are presented as mean ± SEM from 3 replicates. (B) Representative flow cytometry analysis of GsMTx-4 inhibition on Yoda1-induced PS exposure in deoxygenated SS RBCs. Each GsMTx-4 concentration was tested in 3 independent experiments. (C) Dose-response curves of GsMTx-4 inhibition on Yoda1-induced Ca²⁺ influx (black) and PS exposure (red) in deoxygenated SS RBCs. Signals were normalized to the 0 μM GsMTx-4 condition and fitted with the Hill equation (see Methods). (D) IC₅₀ values for GsMTx-4 inhibition of Yoda1-induced Ca²⁺ influx and PS exposure in deoxygenated SS RBCs. Unpaired two-sided Student’s t-test. **P < 0.01. Data were obtained from 3 SCD patients with triplicates for each sample. (E) Comparison of PS^+^ microparticle release from deoxygenated SS RBCs after 2 μM Yoda1 treatment in the absence and presence of 5 μM GsMTx-4. Unpaired two-sided Student’s t-test. ****P < 0.0001. Data were obtained from 5 sickle cell patients with triplicates for each sample. (F) Representative thrombin-mediated fluorogenic results to report prothrombinase activity of SS RBCs under three different conditions: deoxygenation without Yoda1, deoxygenation followed by Yoda1, and deoxygenation followed by Yoda1 and 5 μM GsMTx-4 treatment. (G) Quantification of prothrombinase activity under the conditions in (F), measured as the area under the curve (AUC). One-way ANOVA followed by Tukey’s test. ***P < 0.001, ****P < 0.0001. Data was obtained from 3 SCD patients with duplicates for each sample.

We also found that 5 μM GsMTx4 significantly suppressed Yoda1-induced PS^+^ RMPs release from deoxygenated SS RBCs (Figure S4 and Figure 5E), further supporting the effectiveness of this strategy. Since both PS^+^ RBCs and PS^+^ RMPs contribute to coagulation (van Tits et al., 2009; Whelihan et al., 2016; Whelihan and Mann, 2013; Whelihan et al., 2012), we further assessed the impact of GsMTx4 on thrombin generation using a fluorogenic prothrombinase assay (Evtugina et al., 2023; Smith et al., 1980). 5 μM GsMTx4 largely prevented thrombin generation from Yoda1-stimulated, deoxygenated SS RBCs (Figure 5F-G), demonstrating that partial inhibition of PIEZO1 to break its coupling to TMEM16F could be an attractive approach to reduce SCD-associated thrombotic risk.

### Benzbromarone (Benz) decouples PIEZO1-TMEM16F in deoxygenated SS RBCs

GsMTx4, a peptide derived from tarantula venom (Bae et al., 2011; Gottlieb et al., 2007), has not been approved for human use, presenting a significant obstacle to its clinical translation (Xiao, 2024). To overcome this limitation, we recently identified benzbromarone (Benz), an anti-gout medicine, as a potent PIEZO1 inhibitor that can effectively disrupt PIEZO1-TMEM16F coupling and reduce excessive PS exposure in HX RBCs (Liang et al., 2024). We therefore sought to evaluate Benz’s therapeutic potential in preventing excessive PS exposure in deoxygenated SS RBCs.

Similar to GsMTx4, 5 µM Benz significantly attenuated both the deoxygenation-induced elevation of [Ca^2+^]i (Figure 6A) and PS exposure in SS RBCs under basal conditions (Figure 6B). In deoxygenated SS RBCs, Benz also inhibited 2 µM Yoda1-induced [Ca^2+^]i increase (Figure 6C) and PS exposure (Figure 6D) with IC50 of 2.48 ± 0.41 µM and 0.52 ± 0.01 µM, respectively (Figure 6E-F). Our confocal fluorescence imaging experiments to monitor Yoda1-induced PS exposure confirmed the effectiveness of 5 µM Benz on breaking PIEZO1-TMEM16F coupling and preventing PS exposure in deoxygenated SS RBCs (Figure S5). The divergence of the IC50s of Benz to inhibit Yoda1-induced [Ca^2+^]i and PS exposure (Figure 6E-F) further highlights the feasibility of utilizing this crucial therapeutic window to prevent PS exposure in SS RBCs while still maintaining PIEZO1’s essential physiological functions.

**Figure 6.**
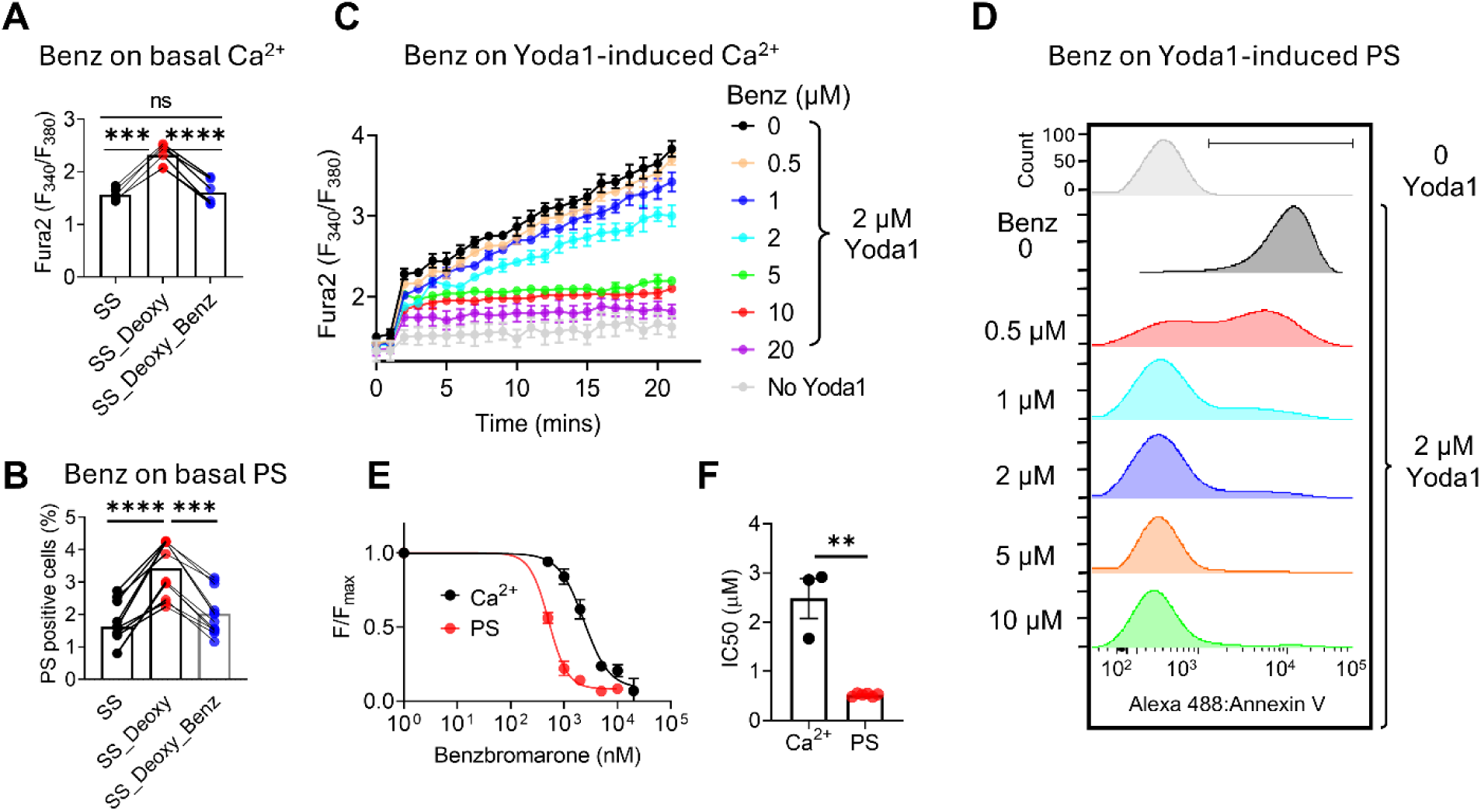
Benz breaks PIEZO1-TMEM16F coupling, reducing Yoda1-induced Ca^2+^ increase and PS exposure in deoxygenated SS RBCs. (A) Basal intracellular Ca²⁺ levels in SS RBCs under normoxic condition and deoxygenated conditions with and without 5 µM benzbromarone (Benz). One-way ANOVA followed by Tukey’s test. ***P < 0.001, ****P < 0.0001, ns: no significance. n = 4 SCD patients. Data are presented as the means of triplicates for each sample. (B) Percentage of basal PS+ SS RBCs under normoxic and deoxygenated conditions with and without 5 µM Benz. One-way ANOVA followed by Tukey’s test. ***P < 0.001, ****P < 0.0001. n = 6 SCD patients. Data are presented as the means of triplicates for each sample. (C) Fura2 measurements of Yoda1-induced Ca²⁺ influx in deoxygenated SS RBCs in response to different concentrations of Benz. n = 3 SCD patients. Data are presented as mean ± SEM from 2–3 replicates. (D) Representative flow cytometry analysis of Benz inhibition on Yoda1-induced PS exposure in deoxygenated SS RBCs. Each Benz concentration was tested in 3 independent experiments. (E) Dose-response curves of Benz inhibition on Yoda1-induced Ca²⁺ influx (black) and PS exposure (red) in deoxygenated SS RBCs. Signals were normalized to the 0 μM Benz condition and fitted using the Hill equation (see Methods). (F) IC₅₀ values for Benz inhibition of Yoda1-induced Ca²⁺ influx and PS exposure in deoxygenated SS RBCs. Unpaired two-sided Student’s t-test. **P < 0.01. n = 3 SCD patients for Ca²⁺ influx and n = 6 for PS exposure. Data are presented as the means of triplicates for each sample. The lines in panels A and B connect data from the same individuals.

### Benz suppresses thrombin generation in SS RBCs

5 µM Benz significantly reduced Yoda1-induced PS^+^ RMPs release by over 70% from deoxygenated SS RBCs (Figure 7A-B). Since PS^+^ RBCs and RMPs contribute to procoagulation, we next evaluated the effectiveness of Benz to inhibit thrombin generation in SS RBCs. In the absence of Yoda1 stimulation, thrombin generation in SS RBCs was significantly increased after deoxygenation (Figure 7C-D), consistent with deoxygenation-induced increase of basal PS exposure and PS⁺ RMPs (Figure 1J-K). Yoda1 stimulation greatly promoted thrombin generation in SS RBCs, and deoxygenation resulted in significantly faster and more thrombin generation (Figure 7C-D). This aligns with increased PS⁺ SS RBCs and RMPs following Yoda1 application (Figure 2). Importantly, 5 µM Benz significantly suppressed Yoda1-induced thrombin generation in deoxygenated SS RBCs, supporting its inhibitory effect on Yoda1-induced PS exposure in SS RBCs (Figure 6D) and PS⁺ RMP release (Figure 7A-B).

**Figure 7.**
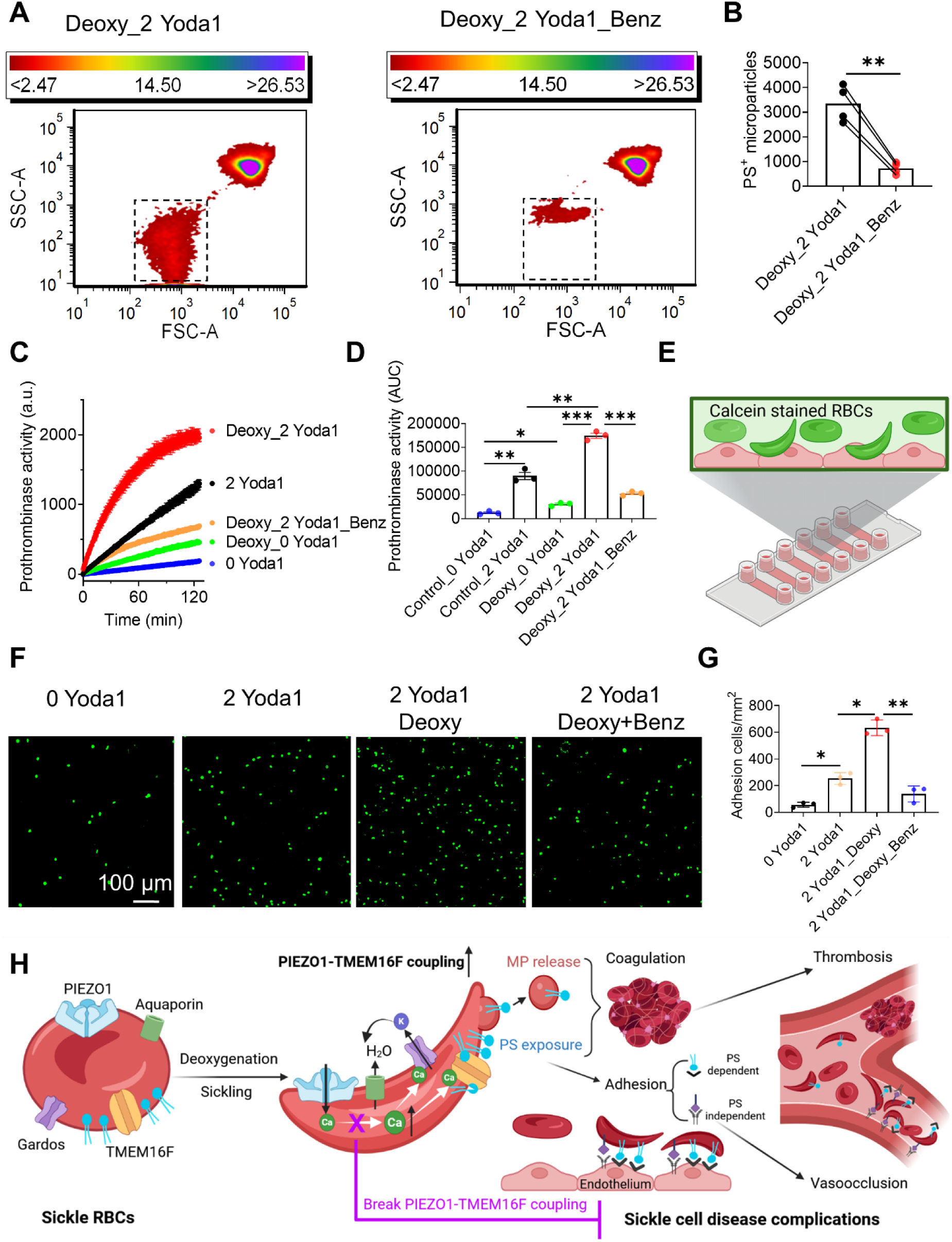
Benz suppresses sickling-induced PS^+^ microparticle release, thrombin generation, and adhesion to endothelial cells. (A) Representative flow cytometry plots of 2 µM Yoda1-induced microparticle release from deoxygenated SS RBCs treated with (right) and without (left) of 5 µM Benz. (B) Comparison of PS+ microparticle release from deoxygenated SS RBCs after 2 μM Yoda1 treatment in the absence and presence of 5 μM Benz. Unpaired two-sided Student’s t-test. **P < 0.01). n=6 SCD patients. Data are presented as the means of triplicates for each sample. (C) Representative thrombin-mediated fluorogenic results to report prothrombinase activity of SS RBCs under five different conditions: normoxic without Yoda1, normoxic with 2 μM Yoda1 stimulation, deoxygenation without Yoda1, deoxygenation followed by Yoda1, and deoxygenation followed by Yoda1 and 5 μM Benz treatment. Data are presented as the means of triplicates for each condition. (D) Quantification of prothrombinase activity under the conditions in (C), measured as the area under the curve (AUC). One-way ANOVA followed by Tukey’s test. *P < 0.05, **P < 0.01, ***P < 0.001. n=3 SCD patients. Data are presented as the means of triplicates for each sample. (E) Schematics of RBCs-endothelium adhesion assay (See methods for details). (F) Representative fluorescence images of the adhesion of calcein-loaded (green) SS RBCs under four different conditions: normoxic without Yoda1 (0 Yoda1), normoxic with 2 μM Yoda1 (2 Yoda1), deoxygenation with 2 μM Yoda1 (2 Yoda1_Deoxy), and deoxygenation followed by 2 μM Yoda1 and 5 μM Benz treatment (2 Yoda1 Deoxy + Benz). (G) Quantification of adhered RBCs to endothelial cells under the conditions in (F). One-way ANOVA followed by Tukey’s test. *P < 0.05, **P < 0.01. n=3 SCD patients. Data are presented as the means of duplicates for each sample. (H) Partial inhibition of PIEZO1 disrupts the functional coupling between PIEZO1 and TMEM16F in SS RBCs, preventing TMEM16F-mediated PS exposure and PS-induced SCD-associated complications including hypercoagulability and vaso-occlusion.

### Benz attenuates SS RBC endothelial adhesion

Abnormal adhesion of sickled SS RBC to the endothelium leads to VOC, a common complication of SCD (Kaul et al., 2009a; Kaul et al., 2009b). A recent study identified the connection between PIEZO1 activation and changes in the adhesive properties of SS RBCs, including BCAM binding affinity (Nader et al., 2023). In addition, it has been shown that PS-exposed SS RBCs are more adhesive to endothelial cells (Setty et al., 2002). Therefore, we investigated whether Benz inhibition of PIEZO-TMEM16F coupling could attenuate SS RBC adhesion to human umbilical vein endothelial cells (HUVECs) *in vitro.* To prevent quantification bias using the traditional bright field microscopy, we developed a fluorescence-based adhesion assay in which SS RBCs were loaded with an inert fluorescence dye calcein (Figure 7E, left). Our analysis shows that Yoda1 stimulation significantly promoted SS RBC adhesion under normoxic conditions (Figure 7F). Deoxygenation further enhanced Yoda1-stimulated EC adhesion, which was effectively prevented by 5 µM Benz (Figure 7F). Our result thus demonstrates the effectiveness of Benz in preventing SS RBCs-EC adhesion.

## Discussion

SCD remains a major global health burden. Current treatments, ranging from hydroxyurea to gene therapy, face limitations related to cost, accessibility, target specificity, and incomplete therapeutic coverage (Barak et al., 2024; Cavazzana et al., 2024; DeMartino et al., 2024; Migotsky et al., 2022; Telen, 2020; Telen et al., 2019). Therefore, there is a critical need for affordable, mechanism-based therapies that can be widely applied, especially in resource-limited settings. Membrane transport proteins, due to their surface accessibility, represent attractive therapeutic targets. The Gardos K⁺ channel, for example, has been extensively explored (Ataga et al., 2008). The Gardos inhibitor Senicapoc improves RBC survival, but fails to reduce VOC in a phase III clinical trial, underscoring the challenges of translating molecular insights into clinical efficacy.

In this study, we identify the PIEZO1–TMEM16F axis as a previously unrecognized but central contributor to the pathophysiology of SCD (Figure 7H). We show that deoxygenation-induced sickling enhances PIEZO1 activation, leading to increased Ca²⁺ influx and activation of TMEM16F, the sole CaPLSase in RBCs (Liang et al., 2024; Yang et al., 2012). TMEM16F-mediated lipid scrambling results in PS exposure, release of PS⁺ RMPs, thrombin generation, and increased adhesion of sickled RBCs to endothelial cells, key contributors to the hypercoagulability and vascular dysfunction in SCD.

Crucially, we demonstrate that partial inhibition of PIEZO1 effectively disrupts this pathogenic mechanotransduction cascade, leading to reduced PS exposure, RMP release, and adhesion of SS RBCs to endothelial cells. Benz, a clinically approved uricosuric agent to treat gout and hyperuricemia, potently inhibits PIEZO1, and thus emerges as a promising candidate for therapeutic repurposing in SCD. Interestingly, 27-40% of SCD patients develop hyperuricemia as a result of hemolysis and impaired renal excretion (Lebensburger et al., 2017), suggesting that Benz could offer dual benefits in this population. Future preclinical and clinical studies are needed to evaluate the efficacy and safety of benzbromarone, given its potential to induce hepatotoxicity in rare cases (Azevedo et al., 2019; Lee et al., 2008). Beyond Benz, our identification of the PIEZO1–TMEM16F axis as a critical mediator of SCD pathology may also stimulate the development of more selective and safer modulators targeting this pathway.

The functional coupling of PIEZO1 and TMEM16F may represent a generalizable mechanotransduction mechanism beyond SCD. Proteomic studies have placed TMEM16F (ANO6) within the PIEZO1 interactome (Koster et al., 2024; Tuomivaara et al., 2024), and we have recently shown this axis is also critical for trophoblast differentiation and placental development (Zhang et al., 2025), suggesting its broad relevance across cell types and various disease conditions.

We also establish that TMEM16F is essential for PS^+^ RMP release from RBCs, a process long known to be triggered by Ca²⁺ (Thangaraju et al., 2020). Using TMEM16F KO RBCs, we confirm that TMEM16F-mediated lipid scrambling is required for PS⁺ RMP generation (Figure 4). Since PS^+^ RMPs contribute to hypercoagulability and vascular damage in SCD and other hemolytic disorders (Garnier et al., 2020; Shet et al., 2003; Van Dreden et al., 2024; Noulsri et al., 2018), targeting the PIEZO1-TMEM16F axis offers a new strategy to mitigate these downstream complications.

Our electrophysiological recordings further reveal that PIEZO1 activity is markedly enhanced in sickled SS RBCs, not only due to altered channel gating, potentially from post-translational modifications (Zhang et al., 2024), but also from increased channel density on the cell surface (Figure 1C–E). PIEZO1 is known to localize preferentially to inward-curving membrane domains in RBCs (Vaisey et al., 2022). During sickling, RMP shedding, which typically involves outward membrane budding, may lead to selective retention of PIEZO1 and a reduction in membrane surface area, collectively increasing channel density on the sickled SS RBC surface. Moreover, the mechanical properties of sickled cells, including increased stiffness (Barabino et al., 2010; Evans et al., 1984; Li et al., 2017), likely promote PIEZO1 activation by reducing its inactivation rate (Ridone et al., 2020; Romero et al., 2019). These biophysical changes, along with biochemical modifications, synergistically amplify PIEZO1-mediated Ca²⁺ influx and downstream TMEM16F activity.

It is important to note that under our pressure-clamp conditions (-60 mmHg), PIEZO1 currents were detectable only at the single-channel level, with stochastic activation in both normal and sickled SS RBCs (Figure 1C-E). Even in sickled RBCs, only 14 out of 31 patches displayed PIEZO1 activity, consistent with prior estimates of ∼100 PIEZO1 channels per RBC (Vaisey et al., 2022). This contrasts with a recent report of robust macroscopic mechanosensitive currents in SS RBCs with slow activation and little inactivation (Romero et al., 2025). The basis for this discrepancy is unclear but may stem from differences in recording conditions, such as mechanical stimulus amplitude, or possibly the presence of an unidentified mechanosensitive channel distinct from PIEZO1. Further studies are needed to determine the molecular identity and contribution of such channels in SS RBCs.

In summary, our findings reveal the PIEZO1-TMEM16F axis as a central mechanistic pathway driving PS exposure, RMP release, and vascular complications in SCD. This work establishes proof-of-concept for targeting this axis therapeutically and identifies Benz as a candidate for repurposing. Future studies aimed at discovering safer and more selective inhibitors, whether small molecules or biologics, will be instrumental in translating this approach into next-generation therapies that address the complex and multifactorial pathophysiology of SCD.

## Methods

### Red blood cell collection

All human studies were approved by the Institutional Review Board at Duke University (Pro00007816, Pro00109511, and Pro00104933). Written informed consent was obtained from patients with SCD and healthy donors (HDs). Murine blood was isolated either from the retro-orbital plexus of the mice anesthetized with isoflurane or from the heart of the euthanized mice following the approved IACUC protocol (#A057-24-02). Both human and murine blood were first collected into tubes containing Acid-citrate-dextrose (ACD) buffer and then centrifuged at 200×g for 12 minutes. After the supernatant was aspirated, the RBC pellets were washed 2-3 times in PBS using the same centrifugation-resuspension protocol. The washed RBC pellets were finally resuspended in the ACD buffer. Purified RBCs were stored at 4℃ and used within 3 days after purification.

### Sex as a biological variable

Our study examined male and female human subjects or mice, and similar findings are reported for both sexes.

### Flow Cytometry

Flow cytometry was performed using a BD FACS Canto flow cytometer to assess PS exposure on RBCs and microparticle release. Packed RBCs were first diluted 1:10 in Hank’s Balanced Salt Solution (HBSS). 2 µL diluted cells were then added to 200 µL of HBSS containing CF488 tagged Annexin V (AnV) at a 1:125 dilution. Yoda1 was added at the desired concentration, and the mixture was incubated for 10 minutes at room temperature before flow cytometry. For drug testing, RBCs were pre-incubated with the test compound before Yoda1 addition, followed by a 10-minute Yoda1 stimulation under the same conditions. Flow cytometry detects microparticle release by measuring the size of particles in a sample using forward scatter (FSC). The samples were analyzed and data were processed with FCS Express software (De Novo).

### Ca²⁺ measurement

Intracellular Ca²⁺ levels were measured using Fura-2 AM. First, 2 µM Fura-2 AM was added to the suspension with 100 µL packed RBCs and 900 µL of HBSS. The cells were incubated at 37°C for 45 minutes to allow dye loading. After incubation, the cells were centrifuged at 200g for 5 minutes to remove unbound dye, resuspended in fresh HBSS, and incubated at 37°C for an additional 10–15 minutes for de-esterification. Fura-2 fluorescence was recorded on a SpectraMax M5 plate reader, with sequential excitation at 340 nm and 380 nm and emission measured at 510 nm. Baseline fluorescence was recorded for 1 minute before the addition of Yoda1 at the desired concentration. Fluorescence readings were taken every 30 seconds for 10–20 minutes, and the results were analyzed using Microsoft Excel.

### Deoxygenation to induce sickling

Sodium metabisulfite was used to induce RBC sickling (Asakura and Mayberry, 1984). To avoid excessive oxidative stress-induced RBC damage (Wairimu et al., 2023), 0.2% instead of the commonly used 2% sodium metabisulfite in HBSS was freshly made and used. Packed RBCs (100 µL) were added to the 1.5 mL of sodium metabisulfite solution in Eppendorf tubes, which were then sealed with pre-warmed wax to prevent gas exchange. The samples were incubated at 37°C for 2 hours to allow irreversible sickling.

### Fluorescence imaging

For RBC imaging, RBCs were seeded on coverslips in buffer (140 mM NaCl, 5 mM KCl, 2 mM MgCl2, 10 mM HEPES, 2 mM CaCl2, pH 7.4). The coverslips were then placed in buffer containing AnV-CF-488A conjugate (1:140 dilution; Biotium, # 29005) on glass slides. Cells were monitored for 50 seconds before the addition of 2 µM Yoda1 to stimulate PIEZO1. A Zeiss 780 inverted confocal microscope was used to simultaneously monitor the morphology and PS exposure of the RBCs in real-time every 5 seconds. A customized MATLAB program (Mathworks) was written to analyze the AnV fluorescent intensity change over time of each cell, as previously described (Le et al., 2019; Liang and Yang, 2021; Zhang et al., 2020).

### RBCs-endothelium adhesion assay

To evaluate the adhesion of RBCs under flow, phosphate-buffered saline (PBS) was prewarmed to 37°C. Yoda1 was prepared at a final concentration of 2 µM in 1 mL HBSS. 10 µL packed RBCs were diluted 1:100 in 1 mL HBSS and incubated with 1 µM Calcein dye for 15 minutes at 37°C to label the cells. The labeled cells were centrifuged at 200g for 5 minutes to remove excess Calcein, and 1 µL of packed RBCs was resuspended in the 2 µM Yoda1 solution prepared earlier, achieving a final RBC dilution of 1:1000. The mixture was incubated at room temperature for 10 minutes. After incubation, the cells were centrifuged again at 200g for 5 minutes to remove excessive Yoda1 and resuspended in 1 mL of prewarmed PBS. To prepare the flow system (ibidi, # 80606), air bubbles were eliminated by running a syringe pump (Chemyx Fusion 200) at a high flow rate (∼5 mL/min). Approximately 0.5 mL of the RBC suspension was then introduced into the flow chamber and incubated for 10 minutes at 37°C. The flow chamber was washed by the syringe pump at a shear stress of 1 dyne/cm² for 5 minutes to remove the non-adherent cells. The number of adhered RBCs was immediately imaged using an Olympus IX83 microscope and quantified by analyzing three representative images captured for each experimental condition using ImageJ. The average number of adhered RBCs across these images was calculated for quantitative analysis.

## Author contributions

Conceptualization: HY, MJT, YZ, PL; Investigation: PL, YSW, KZS, RC; Methodology: PL, YZ, KZS, MD; Software: YZ; Validation: PL; Formal Analysis: PL, YSW, KZS; Resources: HY, GMA, SK, SJF, MD, MJT; Data Curation: PL, YSW, KZS; Writing – Original Draft: HY, PL; Writing – Review & Editing: all authors; Visualization: PL, YSW, KZS; Supervision: HY; Project Administration: HY; Funding Acquisition: HY, PL, YSW.

## Acknowledgments

We thank Dr. Nigel Key lab for their assistance with the fluorogenic prothrombin generation assay. Cartoons were created with BioRender.com. This work was supported by NIH-R35GM153196 (to HY), American Society of Hematology (ASH) Scholar Award (to PL, 2024-2026) and ASH Graduate Hematology Award (to YSW, 2023-2025). Part of this study was reported at the 2024 ASH Annual Meeting.

## Data Sharing Statement

All study data are included in the article and/or SI Appendix. Contract Dr. Huanghe Yang for data and material requests at.

