## Supplemental figures & Methods for "Targeting PIEZO1-TMEM16F Coupling to Mitigate Sickle Cell Disease Complications"

**Affiliations:**
[1]Department of Biochemistry, Duke University School of Medicine, NC 27710, USA
[2]Department of Medicine, Duke University School of Medicine, NC 27710, USA
[3]Department of Surgery, Duke University School of Medicine, NC 27710, USA
[4]Department of Neurobiology, Duke University School of Medicine, NC 27710, USA

*Correspondence:
Huanghe Yang,

Address: Box 3711, DUMC, Durham, NC 27514, USA


**The PDF file includes:**

Supplementary Methods
Supplementary Figure 1 to Figure 5
Captions for Supplementary Videos 1 and 2

### Supplementary Methods

#### Endothelial cell culture

Human umbilical vein endothelial cells (HUVECs) were cultured in EGM-2 BulletKit (Lonza, #CC-3162, authenticated by the Duke Cell Culture Facility) in a humidified incubator at 37°C and 5% $CO_2$-95% air. For the adhesion assay, cells were seeded in ibidi-treat µ-Slide VI 0.4 (Cat.No: 80606) following the manufacturer's instructions. Briefly, 30 µl of $8\times10^5$ cells/ml HUVECs were seeded in each channel. After cell attachment, each reservoir was filled with 60 µl medium. The adhesion assay was performed after a confluent monolayer was formed.

#### Reagents

Yoda1 (Cat. # 21904) and benzbromarone (Cat. # 19768) were purchased from Cayman Chemical Company. GsMTx-4 was purchased from MedChemExpress (Cat. # HY-P1410). Fluorogenic thrombin substrate, Z-Gly-Gly-Arg-AMC was purchased from Bachem, Switzerland (Cat. # 102601-58-1). Bovine Factors Va (BCVA-1110), Xa (BCXA-1060), and prothrombin (BCP-1010) were purchased from PROLYTIX. All other chemicals were obtained from MilliporeSigma.

#### Prothrombinase Assay

To assess prothrombinase activity, 1 µL of packed RBCs was resuspended in 500 µL of assay buffer containing 3 mM $Ca^{2+}$. The cells were then incubated with 2 µM Yoda1 (prepared from a 1 mM stock solution by adding 1 µL to the suspension) for 10 minutes at room temperature. 1 nM Factor Xa and 2 nM Factor Va were added to the reaction mixture containing RBCs (prepared from stock solutions of 100 nM and 200 nM, respectively, by adding 5 µL of each to the 500 µL suspension) and incubated for 2 minutes at 37°C. Next, 50 µL of the reacted solution was transferred into black 96-well plates, and 1.4 µM prothrombin (prepared from a 36 µM stock solution by adding 2 µL to each well) was added to each well. The plate was incubated at 37°C for 2 minutes, followed by the addition of 100 µL of quenching buffer to stop the reaction. The fluorogenic thrombin substrate, Z-Gly-Gly-Arg-AMC, (Bachem, Switzerland) was dissolved in DMSO at a stock concentration of 80 mmol/l. The fluorogenic substrate was then added at a final concentration of 5 µM (0.75 µL to each well). The fluorescence signal was recorded at excitation $\lambda$ = 355 nm and emission $\lambda$ = 460 nm, with readings taken every 15–30 seconds. The plate was shaken before and between each reading to ensure homogeneity.

#### Electrophysiology

Currents were recorded in the cell-attached configuration using an Axopatch 200B amplifier (Molecular Devices) and pClamp software (Molecular Devices). Glass pipettes were fabricated from borosilicate capillaries (Sutter Instruments) and fire-polished with a microforge (Narishige) to achieve a resistance of 10–15 MΩ. We performed recordings on randomly selected locations of SS RBCs, capturing PIEZO1 activity in both non-sickled and sickled cells. For the I–V protocol, currents were elicited using voltage steps from -100 mV to +160 mV in 20 mV increments, with a holding potential of -60 mV. $Cs^+$-based solutions were employed to eliminate contaminating currents from Gardos/IK and other $K^+$ channels. To measure PIEZO1 activity, symmetrical solutions were used in the pipette and the bath, consisting of (in mM): 140 CsCl, 10 HEPES, and

1 $MgCl_2$, adjusted to pH 7.4 with CsOH. Mechanical pressure was applied via a high-speed pressure-clamp system (ALA Scientific Instruments, model HSPC-2-SB), and patches were held at -80 mV. For PIEZO1-TMEM16F coupling experiments, the pipette solution contained (in mM): 140 CsCl, 10 HEPES, and 1 $MgCl_2$, adjusted to pH 7.4 with CsOH. The bath solution was composed of 140 CsCl, 10 HEPES, and either 0 or 2.5 mM $CaCl_2$, as indicated, also adjusted to pH 7.4 with CsOH. Yoda1 was applied extracellularly using a perfusion manifold equipped with a 100-µm tip and eight PE10 tubes, each controlled by a separate valve (ALA-VM8, ALA Scientific Instruments). Only one tube was used at a time for solution application. All experiments were performed at room temperature (22–25°C).

#### Quantification of I-V relation and dose-response to compounds

*I-V* curves were constructed from the steady-state peak currents. Individual *I-V* curves were fitted with a Boltzmann function,

$$I(V) = \frac{I_{max}}{1+e^{\frac{-ZF(V-V0.5)}{RT}}} \quad (1)$$

where $I_{max}$ denotes the fitted value for maximal conductance at a given voltage, $V_{0.5}$ denotes the voltage of half maximal activation of conductance, $z$ denotes the net charge moved across the membrane during the transition from the closed to the open state and $F$ denotes the faraday constant.

Dose response curves were fitted with Hill equation,

$$\frac{F}{F_{max}} = \frac{1}{1+\frac{[IC_{50}]}{[drug]}^{H}} \quad (2)$$

where $F/F_{max}$ denotes the corresponding signal normalized to the max fluorescence signal at given stimuli, [drug] denotes free drug concentrations, $H$ denotes Hill coefficient, and $IC_{50}$ denotes the half-maximal inhibition concentration of individual drug.

#### Statistical analysis

All statistical analyses were performed with Excel or Prism software (GraphPad). Two-tailed Student's t-test was used for single comparisons between two groups (paired or unpaired), and one-way ANOVA following by Tukey's test was used for multiple comparisons. Data were represented as mean ± standard error of the mean (SEM) unless stated otherwise. Symbols *, **, ***, **** and ns denote statistical significance corresponding to p-value <0.05, <0.01, <0.001, <0.0001 and no significance (ns), respectively.

**Supplementary figures**

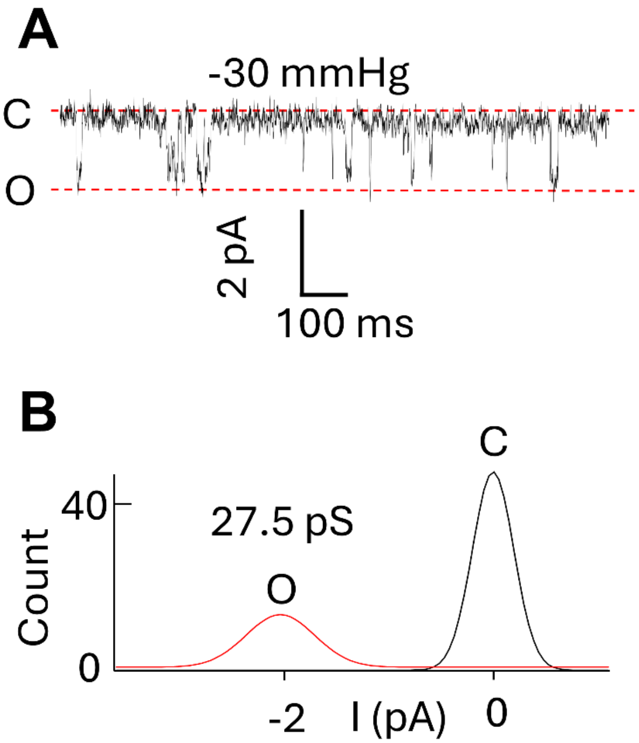


**Figure S1. Single channel characterization of the mechano-sensitive current in sickled SS RBCs.**

(A) Representative single-channel current recorded from sickled SS RBCs under a negative pressure of -30 mmHg applied via pressure clamp. The membrane voltage was held at -80 mV. C: closed state; O: open state.

(B) The amplitude histogram of the mechano-sensitive current shown in A with an average single-channel conductance of 27.5 pS.

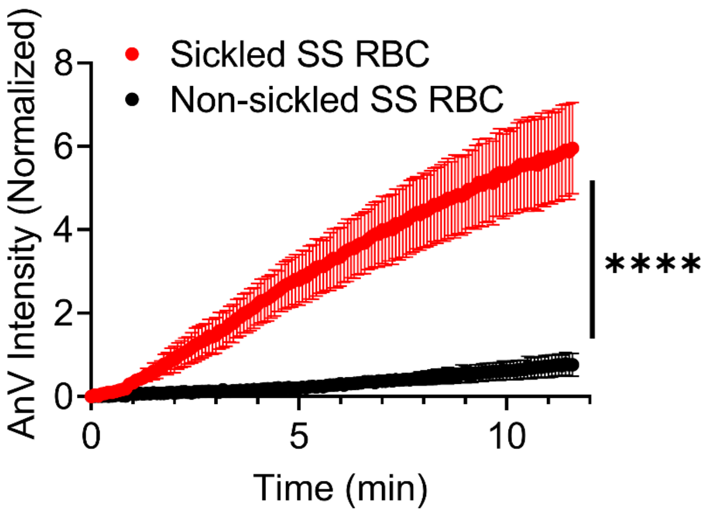


**Figure S2. Sickled-shaped SS RBCs exhibit significantly stronger Yoda1-induced lipid scrambling activity than non-sickled SS RBCs.**

Time-course of 2 µM Yoda1-induced AnV intensity increase in non-sickled and sickled SS RBCs (red star in Fig.2E) under normoxic condition. Statistical comparisons were performed at the final time point using an unpaired two-sided Student's t-test. ****P < 0.0001. Data are presented as the means of triplicates with SEM.

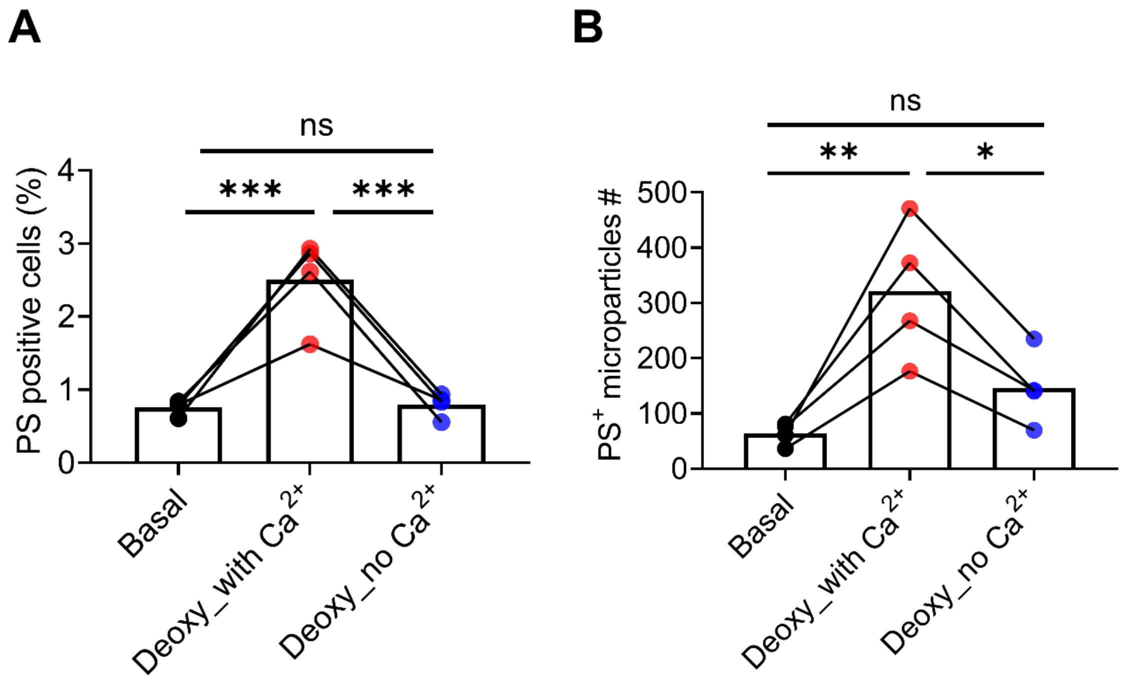


**Figure S3. Deoxygenation-induced PS exposure and PS+ microparticle release are $Ca^{2+}$ dependent.**

(A) Statistical analysis of the percentage of PS+ SS RBCs under the following conditions: normoxic (Basal), deoxygenated with extracellular $Ca^{2+}$ (Deoxy_with $Ca^{2+}$) and deoxygenated without $Ca^{2+}$ (Deoxy_no $Ca^{2+}$). One-way ANOVA followed by Tukey's test. ***$P < 0.001$, ns: no significance. n = 4 SCD patients with triplicates for each sample.

(B) Quantification of $PS^{+}$ microparticle release under the conditions described in (A). One-way ANOVA followed by Tukey's test. *$P < 0.05$, **$P < 0.01$). n=4 SCD patients with triplicates for each sample.

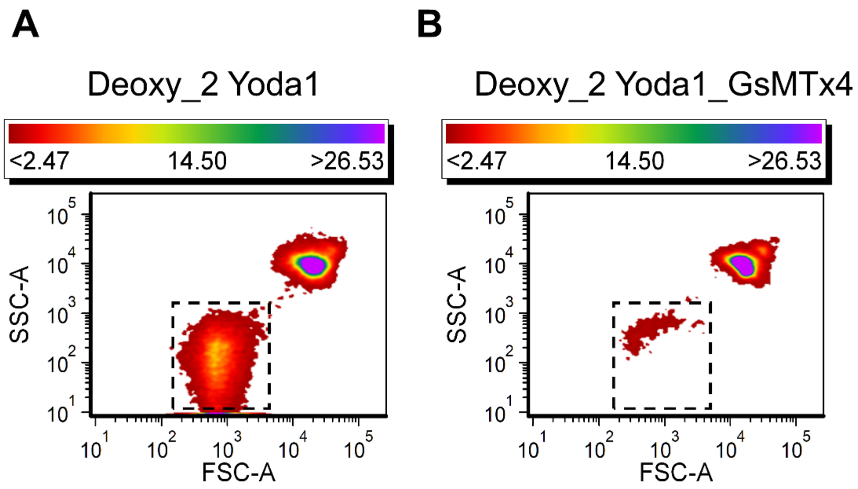


**Figure S4. GsMTx4 suppresses Yoda1-induced microparticle release from deoxygenated SS RBCs.**

(A-B) Representative flow cytometry plots of 2 μM Yoda1-induced microparticle release from deoxygenated SS RBCs treated without (A) and with (B) of 5 μM GsMTx4.

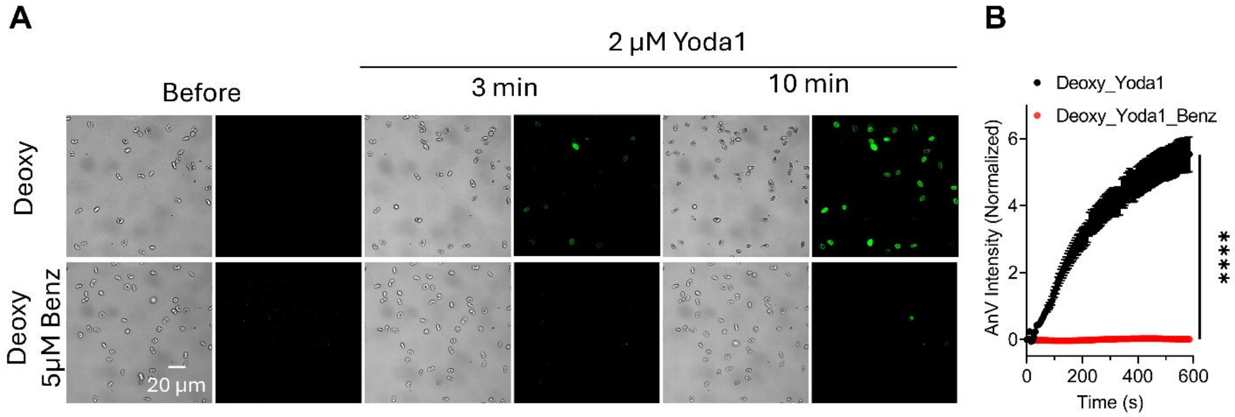


**Figure S5. Benz prevents Yoda1-induced PS exposure in deoxygenated SS RBCs.**

(A) Representative images (bright field on the left and fluorescence AnV on the right) of 2 µM Yoda1-induced PS exposure in deoxygenated SS RBCs treated without (Deoxy_Yoda1) and with (Deoxy_Yoda1_Benz) of 5 µM Benz.

(B) Time-course of Yoda1-induced AnV intensity increase in deoxygenated SS RBCs treated with 2 µM Yoda1 without (black) and with (red) 5 µM Benz. Statistical comparisons were performed at the final time point using an unpaired two-sided Student's t-test. ****$P < 0.001$. Data represents the average of 59 cells (Deoxy_Yoda1) and 57 cells (Deoxy_Yoda1_Benz) from triplicates of one sickle cell patient.

### Supplementary Videos

**Supplementary Video 1.** Simultaneous imaging of RBC morphology and fluorescently tagged Annexin V (AnV-CF488, green) in SS RBCs under normoxic conditions after Yoda1 stimulation (2 μM). Images were acquired every 5 seconds for ~10 minutes. Yoda1 was added at 50 seconds.

**Supplementary Video 2.** Simultaneous imaging of RBC morphology and fluorescently tagged Annexin V (AnV-CF488, green) in deoxygenated sickle RBCs after Yoda1 stimulation (2 μM). Images were acquired every 5 seconds for ~10 minutes. Yoda1 was added at 50 seconds.
